# Degenerate primed randomly amplified polymorphic DNA (DP-RAPD) fingerprinting of bacteriophages of *Vibrio harveyi* from shrimp hatcheries in Southern India

**DOI:** 10.1101/2021.08.10.455891

**Authors:** S. Thiyagarajan, B. Chrisolite, S.V. Alavandi

## Abstract

*Vibrio harveyi* is a significant pathogen of shrimp. Seventy-six bacteriophages infecting luminescent *V. harveyi* were isolated from a total of 194 water samples drawn from various sources of shrimp hatcheries located in South East coast and Andaman island of India. Degenerate primed randomly amplified polymorphic DNA (DP- RAPD) fingerprinting of these bacteriophages was carried out to determine their genetic relatedness. Similarity matrix based on Dice coefficient followed by construction of dendrogram by unweighted pair group method with arithmetic averages (UPGMA) revealed 12 major clusters. One phage was randomly selected from each of these major clusters for transmission electron microscopic observations. Eleven of them had an icosahedral head (46-115 nm) with a long non-contractile tail (132-329 nm), belonging to the *Siphoviridae* family and two phages had a short tail (15-27 nm), belonging to the family *Podoviridae*. The phylogenetic analysis of the phages using DP-RAPD fingerprinting correlated to some extent to the phenotypic nature of the host specifically with regard to sucrose fermentation and source of isolation. However, phages specifically infecting *V. harveyi* and those belonging to different families did not cluster together in the DP-PCR cluster analysis. Hence, the genetic diversity of phages infecting same host with respect to phenotypic difference was revealed by the DP-RAPD applied in this study.

## Introduction

*Vibrio harveyi* has been recognized as a significant pathogen of marine invertebrates, causes extensive mass mortalities of penaeid shrimp larvae in hatcheries resulting in considerable economic losses to shrimp hatchery operators (Karunasagar *et al*. 1994, Austin and Zhang, 2006). Control of LBD has been a challenge to shrimp hatchery operators worldwide. Attempts using antibiotics was reported to be effective in laboratory trials (Baticados *et al*. 1990) while their efficacy in field conditions was reportedly not satisfactory in controlling infections (Moriarty, 1999), in addition to the problems associated with emergence of resistant strains (Karunasagar *et al*.1994).

Bacteriophages, also known as phages are viruses that specifically infect bacteria and lyse them or remain dormant as prophages within their host. Bacteriophages are ubiquitous in nature and found in all habitats that their host bacteria colonize and the hosts are the resource for their proliferation and lifecycle (Weinbauer, 2004). Bacteriophages have been reported to offer an alternative to antibiotics as therapeutic agents in controlling bacterial infections (Defoirdt *et al*. 2011) and the concept of phage therapy has been explored by some investigators in aquaculture (Nakai and Park, 2002, Vinod *et al*. 2006, Karunasagar *et al*. 2007).

Traditional bacteriophage taxonomy is based on their shape, size, morphology, serology, and physiological properties as well as nature of genetic content. According to International Committee on Taxonomy of viruses, bacteriophages have been classified largely based on their morphology and nucleic acid, into one order, Caudovirales, having 14 officially accepted families and at least five other potential families awaiting classification, majority of which comprise tailed phages, belonging to three families, Myoviridae, *Siphoviridae* and *Podoviridae* (Ackermann, 2007, Ackermann, 2009). The size of bacteriophage genomes is reported to be ranging from 15 to 500 kb as seen from the NCBI database. Classical approach to differentiate phages usually involves DNA analysis, biological parameters of phage isolates, analysis of proteins and electron microscopy.

Current approaches include pulsed-field gel electrophoresis (PFGE) and restriction digestion analyses, which allow for the initial classification, while PCR, shotgun cloning and sequence analysis produce more definitive and comprehensive information (Carlson, 2005). Recently, using fluorescence-labeled restriction fragment length polymorphism (fRFLP), it was demonstrated that genetically closely related phages formed clusters and that clustering did not correspond with host range of phages (Merabishvili *et al*. 2007). Restriction digests of genomic DNA and DNA/DNA hybridizations used for examining genetic diversity of related vibriophages (Kellogg *et al*. 1995) and virioplankton of Chesapeake Bay (Wommack *et al*. 1999).

RAPD-PCR using purified DNA has also been used to assess the genetic diversity of vibriophages (Comeau *et al*. 2006, Shivu *et al*. 2007) and phages infecting Escherichia coli (Dini and De Urraza, 2010) and Pseudomonas aeruginosa (Li *et al*. 2010). Our earlier study (Thiyagarajan *et al*. 2011) provided description of three bacteriophages of the family *Siphoviridae* and one phage of *Podoviridae* based on restriction enzyme analysis (REA) and pulsed field gel electrophoresis (PFGE) profile of four phages. The random PCR amplifications of DNA segments using short primers of arbitrary nucleotide sequence have been used to generate specific profiles or genomic fingerprints that are used to compare the genotypic diversity among vibriophages (Shivu *et al*. 2007) and phages infecting Escherichia coli (Dini and De Urraza, 2010) and Pseudomonas aeruginosa (Li *et al*. 2010).

Additionally, low stringency PCR amplification using a degenerate primer in RAPD assay (DP-RAPD) was used to profile algal virus and bacteriophage isolates, encompassing a range of genome sizes, virus families, and hosts (Comeau *et al*. 2004). During our search of bacteriophages as biocontrol agent for luminescent bacterial disease of larval tiger shrimp, we isolated 76 bacteriophages using 27 luminescent *V. harveyi* hosts. The objective of our study was to examine if these phages could be differentiated based on their lystic spectrum or phenotypic nature of the hosts, since the majority of these isolates were from hatcheries located on the coast of Tamil Nadu, South India, and this was evaluated using DP-PCR, cluster analysis and transmission electron microscopic observations.

## Materials and Methods

### Bacteriophages of *Vibrio harveyi*

Seventy-six bacteriophages isolated from shrimp hatcheries were used in this study and were designated as *Vibrio harveyi* phage (VHP 01 to VHP 76; Chrisolite *et al*. 2008). Phage propagation was performed in broth culture of their respective host (Carlson, 2005). Briefly, *V. harveyi* hosts (n=27) were sub-cultured in 5 ml peptone yeast extract sea salt (PYSS) broth (Oakey and Owens, 2000) containing 5 g L^-1^ peptone and 3 g L^-1^ yeast extract dissolved in Macleod’s artificial sea salt water (HiMedia, India). One ml of the culture was inoculated into 50 ml of PYSS broth and incubated at 30 °C for 4-6 h in an orbital shaker at 120 rpm to obtain a cell density of approximately 10^8^ colony forming units per ml (cfu ml^-1^). The bacterial cultures were infected with one ml of respective phage stock and further incubated for about 6-8 h at 30 °C. Phage lysate was prepared by centrifugation of the phage infected broth, twice at 10,400× g for 15 min at 4 °C. The supernatant was filtered through 0.45-μm filter (Millipore, USA). Bacteriophages in the filtrate were concentrated by ultracentrifugation at 200,000 x g for 2 h at 4°C, using SW-41 swinging-bucket rotor (Beckman, CA). Phage pellet was resuspended in sterile phage buffer (0.05M Tris-HCl, 0.02M MgSO4, pH 7.5; (Ghosh *et al*.1989). The suspension was treated with DNase I (1 μg ml^-1^) and RNase A (100 μg ml-1, Merck, India) to degrade nucleic acid residues of the host bacteria. Phage concentrate was purified by ultracentrifugation at 490,000 x g for 18 h at 20°C, in a discontinuous Cesium Chloride (CsCl; SRL, India) density gradient (d = 1.7, 1.5, 1.3 g ml^-1^) prepared in phage buffer (Sambrook and Russell, 2001). The band containing phage particles was drawn from the centrifuge tube using sterile needle fitted to a syringe. CsCl was removed by dialysis at 4°C against phage buffer.

### Extraction of phage Nucleic Acid

Total nucleic acid of bacteriophages was extracted using the protocol described earlier (Santos, 1991) with minor modifications. Briefly, 1 ml of CsCl purified phage suspension previously treated with DNase I and RNase A was mixed with 0.5 ml of TESS buffer (0.1M Tris-HCl pH 8, 0.05M EDTA pH 8, 0.3% SDS and 0.1 M NaCl) and incubated at 56 °C for 5 min. 100μg of Proteinase K (Finnzymes, Finland) was added and further incubated at 56 °C or 30 min and extracted twice with phenol: chloroform: isoamyl alcohol (25: 24: 1). The aqueous phase was transferred into a separate tube and mixed with 0.54 volumes of isopropanol to precipitate the nucleic acid. The pellet was washed with 70% alcohol, briefly air dried and re-suspended in 50 μl of sterile Milli-Q water.

### DP- RAPD Genomic fingerprinting and cluster analysis

The degenerate-primed random amplification of polymorphic DNA was carried out according to a method described previously (Comeau, *et al.,* 2004) using R10D primer sequence 5’-GTCASSWSSW-3’ where S and W represent G/C and A/T, respectively. For optimization of the DP-RAPD, reactions were carried at three different concentrations (50 pM, 100 pM and 200 pM) of R10D primer, incorporation of 3 mM Magnesium chloride instead of 1.5 mM concentration in commercial Taq buffer and a gradient of annealing temperature from 35 to 45°C. The optimized PCR reactions were performed in 25 μL reaction mixes containing 2.5 μL of 10 × Taq buffer with 200 μM of each of the deoxynucleoside triphosphates, 3 mM Magnesium chloride, 200 pM R10D primer, 1.5 U of Taq DNA polymerase (Invitrogen, USA) and 40 ng of template DNA. PCR was performed in a thermocycler (Eppendorf, Germany) under the following thermal cycling conditions: initial denaturation at 95°C for 1.5 min, followed by 40 cycles of denaturation at 95°C for 45 s, annealing at 40°C for 3 min, extension at 72°C for 1 min and final extension at 72°C for 10 min.

DP- RAPD PCR products and 100 bp ladders (Invitrogen, USA) were electrophoresed through 1.5% agarose (Himedia, India) in 0.5 × TBE buffer (45 mM Tris–borate, 1mM EDTA; pH 8.0) at 90 V for 1.5 h. Gels were stained with ethidium bromide (0.5 mg mL-1) and photographed using the Quantity One electrophoresis analysis system (Bio-Rad, CA, USA). Gel images were digitally normalized to a single DNA marker to reduce gel-to-gel banding pattern variability, and cluster analysis was carried out using Molecular Analyst software – Fingerprinting II (Version 3.0, Bio-Rad, CA). The similarity matrix was calculated on the basis of the Dice coefficient, and its corresponding dendrogram was deduced using the unweighted pair group method with arithmetic averages (UPGMA).

### Electron microcopy

A 10μL suspension of purified phages was placed on 200 mesh carbon-coated copper grids and stained with 2% potassium phosphotungstate (pH 7.2) for 20 s. Excess stain was removed immediately by placing the grids on blotting paper. The grids were examined in a Transmission Electron Microscope (Tecnai G2 Spirit Bio-Twin, Eindhoven, The Netherlands).

### Host range

Purified phages (with a titer of about 10^8^ pfu mL-1) were tested by spot assay (Carlson, 2005) to test the spectrum of bactericidal activity against 125 isolates of *V.harveyi* and an isolate each of *Vibrio* species such as V*. logei*, *V. fischeri*, *V. splendidus*, *V. alginolyticus*, *V, paraheamolyticus*, *V. anguillarum*, *V. cholerae (Non- O1)*, *V. fluvialis*, *V. mimicus*, *V. ordalii*, *V. vulnificus and V. metschnikovii*.

## Results and discussion

Bacteriophages are considered to be important components of natural microbial ecosystems. They are ubiquitous and most abundant biological entities on earth and play key roles in regulating the microbial balance in every ecosystem through the lysis of bacterial cells or through horizontal gene transfer (Fuhrman, 1999, Wommack and Colwell, 2000, Williamson *et al*. 2003). Despite the high abundance of phages within aquatic environments, application of molecular genetics tools and microscopy to study viral ecology often requires that virus particles are extracted and concentrated.

In this study, the purification of phage particles isolated from shrimp hatchery environment was done by CsCl gradient ultracentrifugation. The purified phage particles were found as faint whitish bands at CsCl density near to 1.5 gm mL-1. Known tailed phages, belonging to all three families and infecting many different Gram negative and Gram-positive bacteria, are all composed of approximately equal amounts of protein and DNA, giving them a buoyant density in CsCl between 1.45 and 1.52 gm mL-1 (average 1.49; lipid-containing phages average 1.3 g mL-1 (Fraenkel-Conrat, 1985). The titer of phages in the purified stock was estimated to be in the range of 2.8 × 1012 to 8 × 1012 pfu mL-1.

The DNA extraction from the phage concentrates yielded 0.2 to 1μg of DNA. Appropriate dilutions of DNA were made to get uniform template concentration (40 ng). However, recently it was reported that phage suspensions are also suitable to generate reproducible RAPD profiles, bypassing the need for isolating DNA, suggesting that RAPD-PCR could be an easy technique to assess the genetic diversity among phages (Gutierrez *et al*. 2011).

### Genomic fingerprinting of phages by DP-RAPD

DP-RAPD profile of 76 phages showed number of bands ranging from 1 - 12, with amplicon sizes ranging from 126 to 1800 bp. Fifty-six of these phages produced 2-6 bands,15 phages produced 8-12 bands and five phages produced only single band of about 537 bp size. The earlier report (Comeau, *et al.,* 2004) using the same primer profiled algal viruses and bacteriophages encompassing an array of genome sized and virus families and reported that the fingerprints were generally composed of a fewer bands than the larger algal virus fingerprints but there was only a moderate linear relationship between genome size and the number of bands. They reported that closely related viruses such as those infecting Micromonas pusilla generated similar, yet unique patterns within the same family of Phycodnaviridae readily distinguishable from those infecting Chlorella like alga.

Among several PCR based DNA fingerprinting methods used by various earlier workers for molecular typing of bacterial viruses, RAPD uses arbitrary primers to detect changes in the DNA sequence in the genome (Gutierrez *et al*.2011). This approach has been reported to be useful for molecular epidemiological typing as it is relatively fast and easy (Williams *et al*. 1990). As it has been described previously, phages infecting distantly related bacterial hosts typically share little or no nucleotide sequence similarity, while phages infecting a specific bacterial host are more similar (Hatfull, 2008). The degeneracy of the primer used in the present study appears to be useful in random amplification of polymorphic sequences in the smaller genome of these bacteriophages. However, these methods are dependent on laboratory conditions such as template DNA concentrations and PCR conditions (Ellsworth, *et al.,* 1993).

To establish usefulness of DP-RAPD-PCR protocol to type phages, phages infecting *V. harveyi* from the same source were selected to test several experimental conditions in order to generate reproducible RAPD patterns and gain a preliminary insight into the discriminating power of this approach. Since RAPD-PCR reactions are considerably influenced by primers and their concentration (Johansson, *et al.,* 1995), R10 D primer was tested at three different concentrations (50, 100 and 200 pM). Lower primer concentrations produced less defined bands for the primer (data not shown). Furthermore, we found that by increasing the concentration of magnesium chloride, better defined band patterns were obtained. It has been described that Mg2+ions form complexes with dNTPs, primers and template DNA, stimulating the action of DNA polymerase (Pomp and Medrano, 1991). Optimal results were obtained by the addition of 3 mM magnesium and 200 pM primer concentration. The optimum annealing temperature was 40°C as revealed by gradient PCR, which showed clear and distinguishable profile of the phage DNA. The temperature below the optimum resulted more bands, which was difficult to discriminate the bands resulting in a streak. Likewise, the annealing temperature above the optimum produced few or no bands.

### Cluster analysis of phages

Cluster analysis of DP- RAPD profiles separated the 76 phages into 12 groups at 34% hierarchical level (Fig 1). One phage was randomly selected from each of the 12 groups for morphological observations of the phages falling in each of these clusters. Eleven of them had head of 46-115 nm size with a long, noncontractile tail of 132-172 × 7-13 nm, belonging to the *Siphoviridae* family and two phages had head (72-77 nm) with a short tail (15-27 nm), thereby belonging to the *Podoviridae* family (Fig 2, Table 1). Cluster analysis revealed that among the 12 phage clusters, clusters 2, 7, 9, 10, 11 and 12A comprised only 2-3 phages. The members of the family *Siphoviridae* were dispersed in the cluster 1 to 11 and cluster 12A.

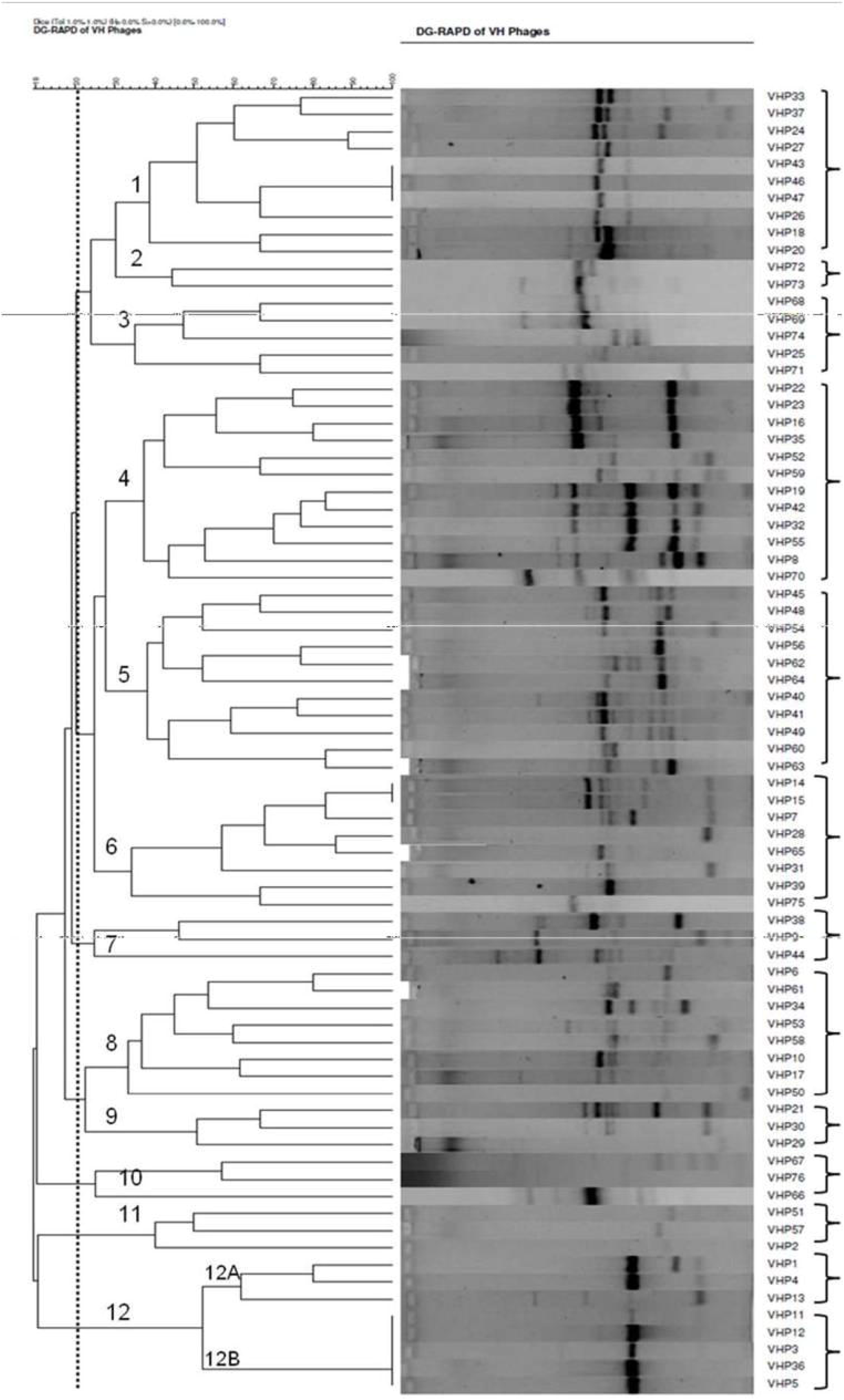

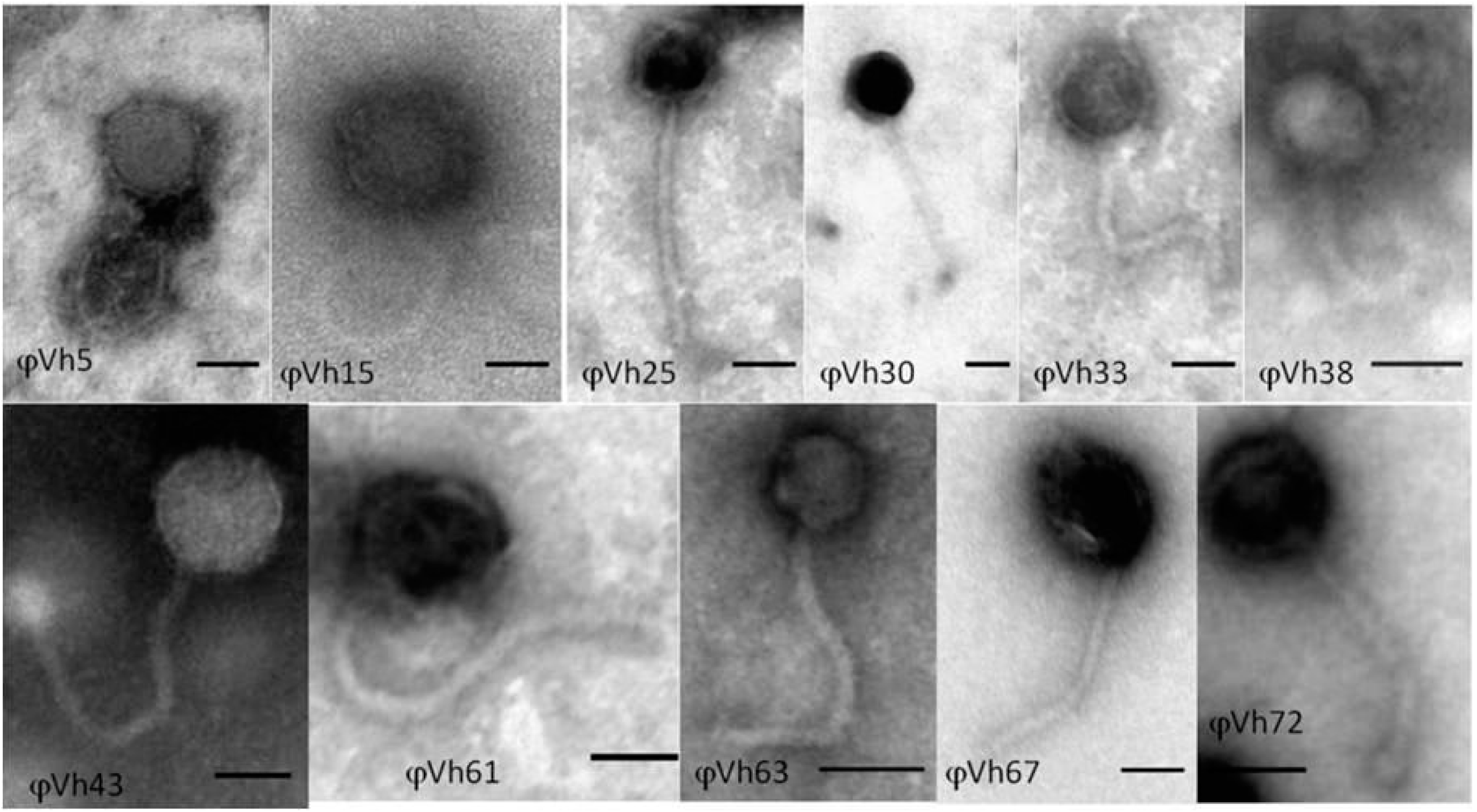

**Table 1.** Characteristics of representative bacteriophages of *V. harveyi* drawn from 12 Clusters

| DG-RAPD Cluster | Phages VHP (n=76) | Lytic activity against 373 phages (average) | Phage Selected for TEM | Head diameter (nm) | Tail length (nm) | Tail width (nm) |
| --- | --- | --- | --- | --- | --- | --- |
| 1 | 33, 37, 24, 27, 43, 46, 47, 26, 18, 20 | 114-221 (176.7) | VHP33<br>VHP43 | 83 ($\pm 6$)<br>87 ($\pm 6$) | 200<br>247 | 12 ($\pm 3$)<br>9 ($\pm 3$) |
| 2 | 72, 73 | 2-41 (21.5) | VHP72 | 69 ($\pm 7$) | 153 | 13 ($\pm 3$) |
| 3 | 68, 69, 74, 25, 71 | 5-41 (16.2) | VHP25 | 73 ($\pm 12$) | 206 | 13 ($\pm 6$) |
| 4 | 22, 23, 16, 35, 52, 59, 19, 42, 32, 55, 08, 70 | 84-189 (137.33) | VHP16 | 84 ($\pm 7$) | 230 | 23 ($\pm 7$) |
| 5 | 45, 48, 54, 56, 62, 64, 40, 41, 49, 60, 63 | 62-224 (139.3) | VHP63 | 47 ($\pm 6$) | 153 | 12 ($\pm 3$) |
| 6 | 14, 15, 07, 28, 65, 31, 39, 75 | 80-250 (157.5) | VHP15 | 108 ($\pm 6$) | 210 | 13 ($\pm 3$) |
| 7 | 38, 09, 44 | 87-94 (90) | VHP38 | 60 ($\pm 6$) | 166 | 10 ($\pm 3$) |
| 8 | 06, 61, 34 53, 58, 10, 17, 50 | 114-220 (160.13) | VHP61 | 76 ($\pm 6$) | 226 | 9 ($\pm 3$) |
| 7 | 38, 09, 44 | 87-94 (90) | VHP38 | 60 ($\pm 6$) | 166 | 10 ($\pm 3$) |
| 8 | 06, 61, 34 53, 58, 10, 17, 50 | 114-220 (160.13) | VHP61 | 76 ($\pm 6$) | 226 | 9 ($\pm 3$) |
| 9 | 21, 30, 29 | 146-232 (190.7) | VHP30 | 66 ($\pm 9$) | 195 | 10 ($\pm 3$) |
| 10 | 67, 76, 66 | 57-70 (69.7) | VHP67 | 96 ($\pm 3$) | 220 | 16 ($\pm 6$) |
| 11 | 51, 57, 02 | 142-175 (159.3) | VHP02 | 60 ($\pm 12$) | 130 | 14 ($\pm 1$) |
| 12A | 01, 04, 13 | 172-218<br>(197.6) | VHP01 | 80 ($\pm 7$) | 170 | 15 ($\pm 1$) |
| 12B | 11, 12, 03, 36,<br>05 | 192-239<br>(210.3) | VHP03* | 72 ($\pm 5$) | 27 | 17($\pm 1$) |
| | | | VHP05* | 77 ($\pm 4$) | 15 | 10 ($\pm 2$) |

\**Podoviridae*

The cluster 12B grouped five phages (VHP3, 5, 11, 12 and 36) in a single clade which comprised the two *podoviruses* (φVH05 in Fig.2). The clusters matched to some extent with the phenotypic nature of the host and the source of isolation. Ten phages in cluster 1 were isolated from hatchery water sample, which were able to ferment sucrose and produce extra-cellular proteases (ECPs) (data not shown). Two phages in cluster 2 were isolated form water samples of Andaman Islands, geographically away from the mainland, and these phages had limited lytic spectrum against *V. harveyi* (Table 1).

Except the phages in cluster 2 and 10, phages in other four clusters showed wide spectrum of lytic activity ranging from 87-239 of the 373 *V. harveyi* isolates. Cluster 3 composed of five phages infecting *V. harveyi* that were positive for sucrose fermentation and negative for ECP production. Clusters 4, 5 and 6 consisting of 31 phages virulent to *V. harveyi* isolates showed variable sucrose fermentation activity and negative for production of ECP. The phages of cluster 1 to 11 were isolated during larval production cycle affected by luminescent bacterial disease (LBD) except the phages from Andaman source. Fifty two of the 76 phages in the present study were able to lyse other closely related vibrios *like V. alginolyticus*, *V. parahaemolyticus and V. logei*.

Fifteen phages, VHP06, VHP09, VHP11, VHP17, VHP18, VHP24, VHP25, VHP28, VHP29, VHP38, VHP39, VHP43, VHP46, VHP52 and VHP54 were specific to *V. harveyi*. However, these phages did not cluster in the DP-PCR cluster analysis. These observations are in conformity with the earlier findings (Gutierrez *et al*. 2011), wherein, their study indicated that genetically different phages infecting same host clustered together based on RAPD fingerprinting. Similar observations were earlier reported and clustering of genomically and serologically closely related phages based on the fingerprints obtained for Caudovirales using fRFLP, wherein, the host range was reported to be not completely in correspondence with genotype (Merabishvili *et al*. 2007).

## Conclusions

DP-RAPD of bacteriophages of *V. harveyi* was found to be suitable to assess the genetic diversity among those infecting *V.harveyi*. The phylogenetic analysis based on DP- RAPD fingerprints separated 76 phages of *V.harveyi* into 12 different clusters which to some extent correlated with phenotypic nature of host bacteria specifically with regard to sucrose fermentation and source of isolation.

## Acknowledgements

Authors are thankful to the Indian Council of Agricultural Research (ICAR), New Delhi, for the financial assistance in carrying out this study (F. No. 4. 57/2004-ASR-I). We also thank Dr. A. G. Ponniah, Former director and Dr. K. K. Vijayan, Director, ICAR - Central Institute of Brackishwater Aquaculture (CIBA) for the critical evaluation of this work.

## Notes

### Competing Interest Statement

The authors have declared no competing interest.

## References

Ackermann HW (2009) Phage classification and characterization. In: Clokie MRJ, Kropinski AM (eds.) Bacteriophages,: Methods and Protocols, Isolation, Characterization and Interactions Vol. I. Humana Press, Clifton, NJ. pp. 127–140.

Ackermann HW (2007) 5500 Phages examined in the electron microscope. Arch Virol 152: 227–243.

Austin B & Zhang XH (2006) *Vibrio harveyi:* a significant pathogen of marine vertebrates and invertebrates. Lett Appl Microbiol 43: 119–124.

Baticados MCL, Lavilla-Pitogo CR, Cruz-Lacierda ER, Pena LD, Sunaz NA (1990) Studies on the chemical control of luminous bacteria *Vibrio harveyi* and *V. splendidus* isolated from diseased *Penaeus monodon* larvae and rearing water. Dis Aquat Organ 9: 133–139.

Carlson K (2005) Working with Bacteriophages: Common techniques and methodological approaches. In: Kutter E, Sulakvelidze A (eds) Bacteriophages: Biology and Applications, CRC Press, Boca Raton. pp. 437–494.

Chrisolite B, Thiyagarajan S, Alavandi SV, Abhilash EC, Kalaimani N, Vijayan KK, Santiago TC (2008) Distribution of luminescent *Vibrio harveyi* and their bacteriophages in a commercial shrimp hatchery in South India. Aquaculture 275: 13–19.

Comeau AM, Short S, Suttle CA (2004) The use of degenerate-primed random amplification of polymorphic DNA (DP-RAPD) for strain-typing and inferring the genetic similarity among closely related viruses. J Virol Meth 118: 95–100.

Comeau AM, Chan AM, Suttle CA (2006) Genetic richness of vibriophages isolated in a coastal environment. Environ Microbiol 8: 1164–1176.

Defoirdt T, Sorgeloos P, Bossier P (2011) Alternatives to antibiotics for the control of bacterial disease in aquaculture. Cur Opinion Microbiol 14: 251–258.

Dini C, De Urraza PJ (2010) Isolation and selection of coliphages as potential biocontrol agents of enterohemorrhagic and Shiga toxin producing *E. coli* (EHEC and STEC) in cattle. J Appl Microbiol 109: 873–887.

Ellsworth DL, Rittenhouse KD & Honeycutt RL (1993) Artifactual variation in randomly amplified polymorphic DNA banding patterns. Biotechniques 14: 214.

Fraenkel-Conrat H (1985) The Viruses: Catalogue, characterization and classification. Plenum Press.

Fuhrman JA (1999) Marine viruses and their biogeochemical and ecological effects. Nature 399: 541–548.

Ghosh AN, Ansari MQ, Datta GC (1989) Isolation and morphological characterization of E1 Tor cholera phages. J Gen Virol 70: 2241–2243.

Gutierrez D, Martin-Platero AM, Ana Rodriguez A, Martinez-Bueno M, Garcia P, Martinez B (2011) Typing of bacteriophages by randomly amplifed polymorphic DNA (RAPD)-PCR to assess genetic diversity. FEMS Microbiol Lett 322: 90–97.

Hatfull GF (2008) Bacteriophage genomics. Cur Opinion Microbiol 11: 447–453.

Johansson ML, Quednau M, Molin G, Ahrne S (1995) Randomly amplified polymorphic DNA (RAPD) for rapid typing of *Lactobacillus plantarum* strains. Lett Appl Microbiol 21: 155–159.

Karunasagar I, Pai R, Malathi GR, Karunasagar I (1994) Mass mortality of *Penaeus monodon* larvae due to antibiotic-resistant *Vibrio harveyi* infection. Aquaculture 128: 203–209.

Karunasagar I, Shivu MM, Girisha SK, Krohne G, Karunasagar I (2007) Biocontrol of pathogens in shrimp hatcheries using bacteriophages. Aquaculture 268: 288–292.

Kellogg CA, Rose JB, Jiang SC, Thurmond JM, Paul JH (1995) Genetic diversity of related vibriophages isolated from marine environments around Florida and Hawaii, USA. Mar Ecol Progr Ser 120: 89–98.

Li LLL, Yang HYH, Lin SLS, Jia SJS (2010) Classification of 17 newly isolated virulent bacteriophages of *Pseudomonas aeruginosa*. Canadian J Microbiol 56: 925–933.

Merabishvili M, Verhelst R, Glonti T et al (2007) Digitized fluorescent RFLP analysis (fRFLP) as a universal method for comparing genomes of culturable dsDNA viruses: application to bacteriophages. Res Microbiol 158:572–581.

Moriarty DJW (2000) Disease control in shrimp aquaculture with probiotic bacteria. In: Bell CR, Brylinsky M, Johanson-Green PC (eds.), Atlantic Canada Society for Microbial Ecology, Halifax, Canada. pp. 237–244.

Nakai T, Park SC (2002) Bacteriophage therapy of infectious disease in aquaculture. Res Microbiol 153: 13–18.

Oakey HJ, Owens L (2000) A new bacteriophage, VHML, isolated from a toxin-producing strain of *Vibrio harveyi* in tropical Australia. J Applied Microbiol 89: 702–709.

Pomp D, Medrano JF (1991) Organic solvents as facilitators of polymerase chain reaction. BioTechniques 10: 58–59.

Sambrook J, Russell DW (2001) Molecular Cloning: A Laboratory Manual. CSHL press.

Santos MA (1991) An improved method for the small scale preparation of bacteriophage DNA based on phage precipitation by zinc chloride. Nucleic Acids Res 19: 5442–5442.

Shivu MM, Rajeeva BC, Girisha SK, Karunasagar I, Krohne G, Karunasagar I (2007) Molecular characterization of Vibrio harveyi bacteriophages isolated from aquaculture environments along the coast of India. Environ Microbiol 9: 322–331.

Thiyagarajan S, Chrisolite B, Alavandi SV, Poornima M, Kalaimani N, Santiago TC (2011) Characterization of four lytic transducing bacteriophages of luminescent *Vibrio harveyi* isolated from shrimp (*Penaeus monodon*) hatcheries. FEMS Microbiol Lett 325: 85–91.

Vinod MG, Shivu MM, Umesha KR, Rajeeva BC, Krohne G, Karunasagar I, Karunasagar I (2006) Isolation of *Vibrio harveyi* bacteriophage with a potential for biocontrol of luminous vibriosis in hatchery environments. Aquaculture 255: 117–124.

Weinbauer MG (2004) Ecology of prokaryotic viruses. FEMS Microbiol Rev 28: 127–181.

Williams JGK, Kubelik AR, Livak KJ, Rafalski JA, Tingey SV (1990) DNA polymorphisms amplified by arbitrary primers are useful as genetic markers. Nucleic Acids Res 18: 6531–6535.

Williamson KE, Wommack KE, Radosevich M (2003) Sampling natural viral communities from soil for culture-independent analyses. Appl Environ Microbiol 69: 6628–6633.

Wommack KE, Colwell RR (2000) Virioplankton: viruses in aquatic ecosystems. Microbiol Mol Biol Rev 64: 69–114.

Wommack KE, Ravel J, Hill RT, Colwell RR (1999) Hybridization analysis of Chesapeake Bay virioplankton. Appl Environ Microbiol 65: 241–250.

